# Structural Basis for Dual Peptidoglycan Hydrolysis by an *E. faecium Minhovirus* Tail Spike Lysin

**DOI:** 10.64898/2026.03.11.711020

**Authors:** S Mesnage, J Wilson, Y Zhang, A Alrafaie, M Robertson, EM Smith, CA Evans, J Li, JB Rafferty, GP Stafford

## Abstract

Bacteriophages rely on breaching the bacterial cell wall as the first step of infection. We characterise ORF11, a putative tail-spike lysin from the 19 kbp *Minhovirus* SHEF14, a podovirus infecting *Enterococcus faecium*. The ORF11 2.5 Å crystal structure reveals that ORF11 comprises four domains: a predicted glycosyl hydrolase (D1) a cysteine, histidine-dependent amidohydrolase/peptidase (CHAP, D4), separated by a helical linker (D2) and a CHAP-like domain (D3). The crystal form reveals a dimeric assembly with the predicted glycosyl hydrolase (D1) and CHAP peptidase (D4) domains located adjacent to each other (but at opposite ends of the dimer) within one subunit and separated by domains D2 and D3-positioned at the dimer interface. The presence of the D3 CHAP-like domain is unusual among related bacteriophage tail-lysins from related small podoviruses but conserved in *E. faecium Minhoviruses*. Biochemical assays using recombinant ORF11 and LC-MS/MS confirmed dual peptidoglycan-degrading activity. ORF11 functions as both an *N*-acetylglucosaminidase and a D,D-endopeptidase, cleaving the bond between the D-alanine in position-4 and the D-aspartate residue at the end of the side chain. Together, these results provide the first structural description of an *Enterococcus* podovirus tail-spike lysin and demonstrate bifunctional enzymatic activity. This dual action likely facilitates initial surface recognition and localised peptidoglycan degradation during infection of *E. faecium*, offering new insights into how small-genome phages target this clinically significant antimicrobial-resistant pathogen.

## Introduction

*Enterococcus faecium* is a monoderm (Gram-positive) bacterium commonly found in the gastrointestinal tracts of humans and animals and is frequently isolated from hospital settings[1–3]. It is associated with a wide range of infections, including urinary tract infection, endocarditis, bacteraemia, wound and intravenous line infections, and is often recovered from diabetic foot ulcers [3, 4]. Enterococci exhibit a high level of antibiotic resistance and can acquire new resistance determinants, severely limiting the effectiveness of conventional antibiotic treatments[5]. *E. faecium* was the first species reported to acquire vancomycin resistance [6] giving rise to Vancomycin or Glycopeptide Resistant Enterococci (VRE/GRE). These strains are notoriously difficult to eradicate and present a major clinical challenge [7].

A notable feature of the genus *Enterococcus* is their resilience and adaptability, which enables them to persist in high numbers across diverse environments, including wastewater [1]. Unsurprisingly, their bacteriophage predators are also widespread, and phage targeting *Enterococcus* species are increasingly recognised as potential alternatives to antibiotics [8]. A growing diversity of bacteriophages that kill enterococci has now been identified, spanning non-contractile siphoviruses (e.g. *Efquatrovirus*, *Aramisvirus, Saphexavirus*), contractile myoviruses (e.g. *Schiekvirus*, *Kochikohdavirus*) and small podoviruses (e.g. *Salasmavirus*, *Copernicusvirus, Minhovirus*) [9–13]. We recently reported the isolation of SHEF14, a *Minhovirus* infecting *E. faecium* that has a 19 kbp genome, 22 Open-reading frames, a small 54 ± 2 nm head and a narrow host-range [13]. Although other *Minhoviruses* infecting enterococci have been described [14, 15], little is known about how they recognise or infect their host. For other enterococcal bacteriophages, including the *Schiekvirus* SHEF13, host specificity is partly determined by the surface Extracellular Polysaccharide Antigen (EPA) [13, 14], comprising a core repeating unit made of largely rhamnose, and a highly variable portion referred to as EPA decorations [16, 17].

Short-tailed viruses (podoviruses) possess compact DNA genomes sometimes as small as 18 kbp and infect both diderm and monoderm bacteria. Well characterized examples include P22 (*Salmonella typhimurium* [18]);T7 (*E. coli* [19]), Phage C1 (*Streptococcus pneumoniae* [20]) or phage Andhra (*Staphylococcus epidermidis* [21]). These phages typically feature an icosahedral capsid with a short tail, along with a baseplate and tail spike that mediate host recognition and cell wall penetration. Proteins within these structures commonly combine Receptor Binding Protein domains (RBPs) as well as putative extracellular lytic activities such as endopeptidases or glycosyl hydrolases [22]. Within the podoviruses, the picoviruses have small genomes (17-20 kbp) and infect monoderm bacteria. Representatives include phage P68 [23], phi29 [24] and Andhra [25], which are genomically and morphologically different from larger T7-like podoviruses [26]. Cryo-EM reconstructions of several picoviruses have revealed the organisation of their baseplate and tail spike complexes [23, 25, 27] and have suggested locations of lytic domains thought to facilitate cell wall degradation. However, despite these structural insights, only a limited number of these tail or spike associated enzymatic activities have been biochemically characterised.

In this study, we present the first structural and biochemical characterisation of a predicted tail-spike protein, ORF11 from the *minhovirus* bacteriophage SHEF14, and explore the implications of these findings for *Enterococcus* phage biology.

## Results

### Crystal structure of SHEF14 ORF11

SHEF14 is a *Minhovirus* with podovirus morphology recently isolated using *E. faecium* VRE strain E1071 as a host [13]. Its genome encodes 22 ORFs, 10 of which have no assigned function. ORF11 encodes a predicted tail-associated peptidoglycan hydrolase (671aa, 72.1 kDa). The gene encoding ORF 11 is positioned upstream of genes encoding a predicted holin-endolysin pair and is annotated as an NlpC/P60/CHAP peptidase (Fig. 1A, green). To elucidate the structure of the predicted tail-spike lysin ORF11, we produced recombinant ORF11 with a C-terminal His-Tag in *E. coli*.

**Fig 1.**
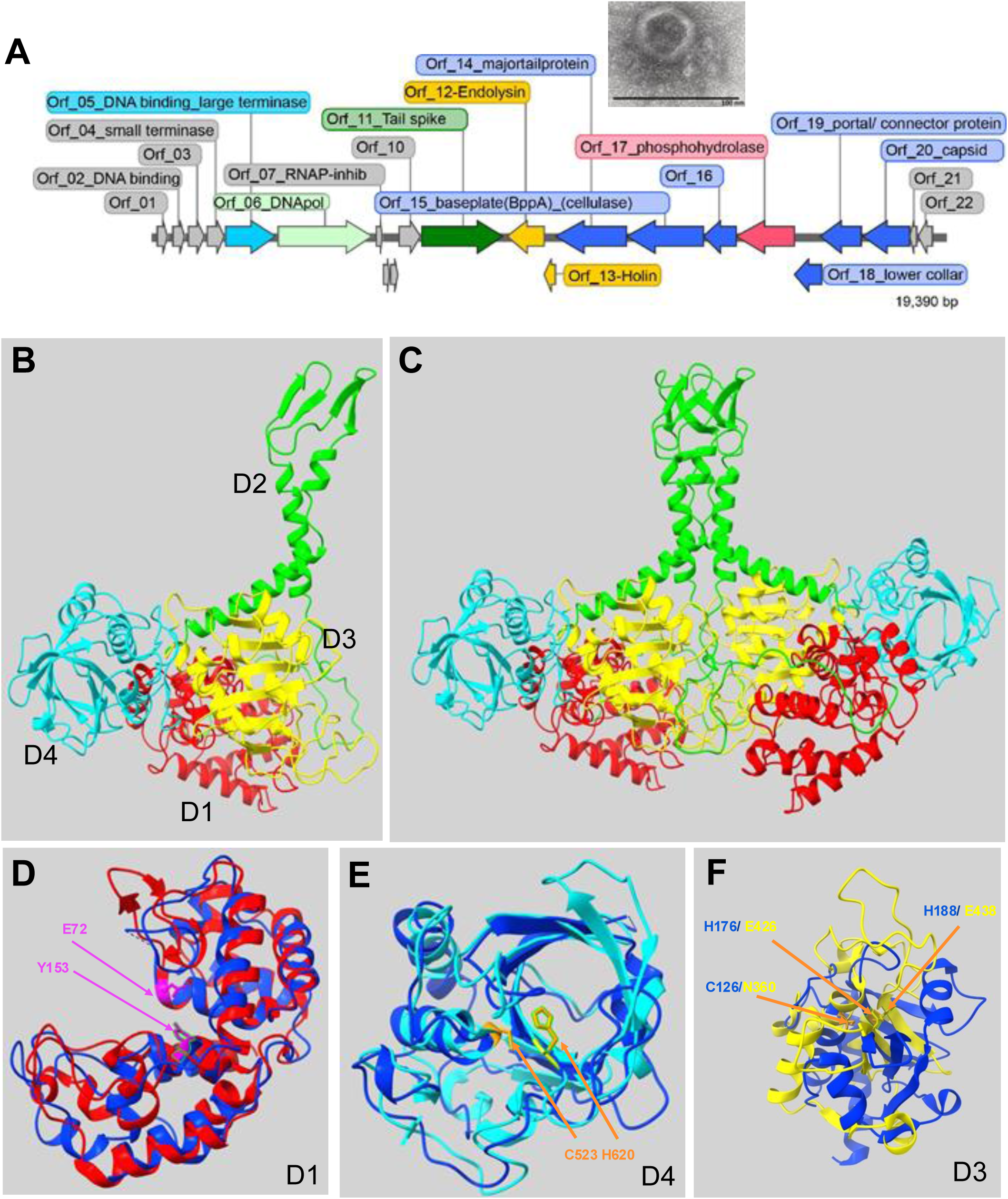
Schematic representation of SHEF14 genome and ORF11 crystal structure. (**A**) Genome annotation of phage SHEF14. Arrows indicate the direction of transcription, and the predicted functions of the ORFs are shown around the genome. The inset shows a TEM image of SHEF14 (**B**) Crystal structure overview of ORF11 monomer with the GH domain (D1, red), CHAP domain (D4, cyan), Linker domain (D2,green) and D3 domain in yellow. (**C**) Crystal structure overview of ORF11 dimer with colours as in (**B**). Structural alignment of ORF11 domains with PlyCA (4f88) homologue is shown for D1 with the GH domain of PlyC; Predicted active site residues are indicated in magenta (**D**), D4 aligned with the CHAP domain of PlyCA (**E**) with active site residues highlighted in orange. The D3 is also shown aligned with the CHAP domain of an endopeptidase from *Anabaena variabilis* (2HBW) (**F)** and locations of equivalent/missing active site residues indicated in blue/yellow.

The ORF11 protein was purified using two successive chromatographic steps (Fig. S1) with gel filtration indicating that it was present as a mixture of dimeric and monomeric forms (most abundant). The protein was successfully crystallised, and X-ray diffraction data were collected. Initial attempts to solve the structure by molecular replacement were unsuccessful even when using AlphaFold models and native, non-derivatized diffraction data. We therefore used selenomethionine-based single wavelength anomalous dispersion (Se-Met SAD) phasing, which enabled calculation of interpretable electron density maps and subsequent model building to 2.5 Å (PDB: 9T7R).

The crystal structure revealed a dimeric unit, with the overall architecture within each monomer consisting of three globular domains (D1, D3, and D4) while D2 adopts an elongated, predominantly alpha helical structure that projects away from the plane of the other domains and ending in a small four-stranded beta sheet (Fig. 1B). The D2 domain forms an extended coiled-coil interface with the corresponding domain in the other monomer involving both the alpha helices and the beta strands which form into a small beta barrel. Outside of the D2 interface, the dimer interface is dominated by interactions between a long loop region leading to D2 with an extended cleft in D3 (details in Fig. S2B,C).

Examination of the ORF11 domain structures, combined with Foldseek and DALI domain comparison searches, revealed that all the three globular domains (D1, D3, and D4) share structural homology with domains of the phage C1 endolysin PlyCA (4f88, [28]) (D1/ D4, Fig. 1D, E) and an endopeptidase from *Anabaena variabilis* (D3, 2HBW,[29]). Domain D1 displays strong structural homology with the PlyCA glycosyl hydrolase domain (GH) (Fig. 1D, Foldseek: prob 1, RMSD 4.6 Å across 217 Cα atoms, TM 0.77) as well as the tail lysin from *Staphylococcus* phages Andhra [25] and P68 [23]. The D4 domain structurally aligns with the PlyCA CHAP domain (Fig 1E, Foldseek: prob 1, RMSD 3.96 Å across 165 residues TM 0.81). In both cases, despite low overall sequence identity (∼16%) between ORF11 and PlyCA, the key catalytic residues are conserved in position: Glu72 and Tyr153 in D1 correspond to the glycosyl-hydrolase active site, and Cys523 and His620 in D4 form the expected CHAP-domain catalytic dyad. This conservation, together with the structural similarity, strongly supports the assignment of D1 and D4 as the enzymatic centres of ORF11. [30].

In contrast, domain D3 which is positioned between domains D1 and D4, does not share homology with PlyCA, but has structural homology with a cell wall endopeptidase from *Anabaena variabilis* (Fig. 1F, 2HBW, [29]) (Foldseek prob 0.91, RMSD 4.11 Å for 139aa, TM 0.66). However, unlike the D1 and D4 domains, it lacks the equivalent CHAP/ papain protease catalytic residues present in this family of proteins-in this case it has no equivalent for C126, H176 or H188 (D3 has an Asparagine, and two glutamines instead-Fig. 1F), indicating that it is unlikely to possess enzymatic activity. Given its position within the overall architecture, D3 may instead contribute to substrate engagement, for example, by coordinating peptidoglycan stem-peptide residues to expand the interaction surface or to orient the adjacent catalytic domains more effectively. This suggests a potential structural or scaffolding/dimerisation role for D3 within the tail-spike assembly.

The individual GH (D1) and CHAP (D4) domains of ORF11 show a strong structural homology to domains in PlyCA as well as counterpart Andhra_Gp10 and P68_ORF11, despite low sequence identity (approx. 20%). However, the overall protein architecture is different to SHEF14 (Fig S2). As shown schematically in Fig. 2A, in SHEF14 ORF11, while the two catalytic domains reside together in 3D space within each dimer they are separated by interacting D3 CHAP-like domain and the extended helical D2 domain, whereas within PlyCA(4f88), Andhra_Gp10(8EJ5_A) and P68_ORF11(AlphaFold) the GH and CHAP domains are separated by only a short helical linker (Fig. 2A). Furthermore, phage SHEF14 is a small podovirus within the genus *Minhovirus*: one of three ‘podovirus’ phage genera that infect enterococci-alongside the *Copernicusvirus* and *Salasmavirus* [12]. The *Minhovirus* and *Copernicusvirus* share a similar genome organisation with nucleotide homology in several ORFs of 25-68% sequence identity (Fig. 2B), but the *Copernicusvirus* ORF11 homologue also lacks the D2 and D3 domains (Fig. 2B). In contrast, the *Salasmavirus* phage Athos [12] contains no obvious sequence or structural homologue of ORF11 (not shown) and only 20-25% nucleotide identity across the entire genome. It contains a gene encoding a protein (orf20, genbank: LR990834.1) structurally homologous to the tail-tip of *Bacillus* phage phi29 (gp13) that contains lysozyme and M23 peptidase domains-i.e. analogous function but no homology (Fig S3). These data indicate that *Enterococcus* podophage contain a diverse set of dual-function tail-tip/spike lysins to degrade the cell wall of this key pathogen.

**Fig 2.**
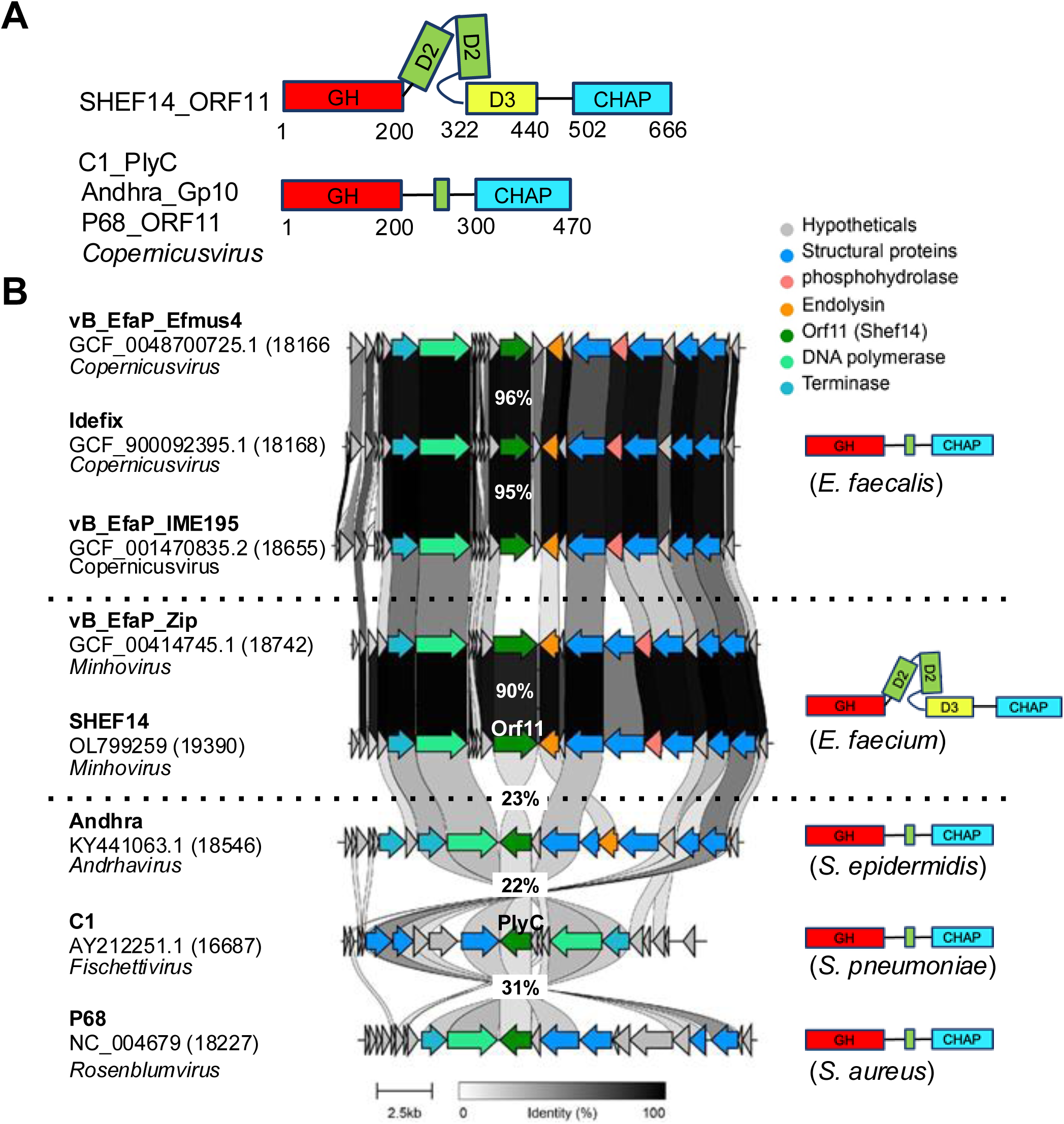
Domain organisation of ORF11 and comparison to phage homologs. (**A**) Schematic organisation of SHEF14 ORF11 and functional homologs in phage C1 (PlyC) and Andhra (Gp10)using same colours as in Fig. 1: GH domain (D1) in red, linker (D2) in green, CHAP homolog (D3) in yellow and CHAP (D4) in cyan. (**B**) Multiple genome sequence alignment. ORF11 is highlighted in dark green. Accession numbers and host species are shown within the diagram. Percentages refer to the amino acid identity compared to ORF11 or PlyC.

Collectively, these findings suggest divergent evolutionary pathways for tail spike proteins among podoviruses infecting monoderm bacteria, with ORF11 and related *E. faecium* phages adopting an architecture distinct from their counterparts in other viruses (Fig. 2).

### SHEF14 ORF11 displays *N*-acetylglucosaminidase and D,D-endopeptidase activities

While PlyCA was proposed to display both glycosyl hydrolase and peptidase activity, the specific bonds cleaved by this enzyme were not identified [31]. To characterise the scissile bonds targeted by ORF11, cell walls were extracted from various *E. faecium* strains [32] to identify the best substrate for this enzyme. The Enterococcal Polysaccharide Antigen (EPA) is one of the surface components critical for phage recognition and can be critical for peptidoglycan cleavage by hydrolases [33]. We therefore tested the activity of purified recombinant His-tagged ORF11 against cell walls from strains belonging to each of the four main EPA variants described in *E. faecium* (E1636, EPA type 1; E1071, type 2; E1162, type 3; and Aus0004, type 4) [17]. Although SHEF14 only infects *E. faecium* E1071, the HPLC muropeptide profiles revealed that ORF11 can solubilise peptidoglycan fragments from all EPA-type strains, with the best activity against Aus0004 cell walls (Fig. S4). This shows that enzymatic activity is not dependent on a specific EPA type, rather it is the conserved peptidoglycan of *E. faecium* that is the target. Based on these preliminary results, we decided to use cell walls from *E. faecium* D344S as this has been previously characterised in detail [32].

To characterise the enzymatic activities of ORF11, *E. faecium* D344S cell walls were digested with mutanolysin alone (a commercially available glycosyl hydrolase cleaving peptidoglycan), ORF11 alone or mutanolysin followed by ORF11 (Fig. 3). The resulting muropeptide profiles showed that ORF11 produced predominantly peptidoglycan fragments with short retention times, consistent with monomeric disaccharide-peptides (Fig. 3A). Analysis of deconvoluted LC-MS data revealed that only 0.28% of muropeptides solubilised by ORF11 were crosslinked, compared to 27.23% in mutanolysin-digested material (an expected result since mutanolysin cannot cleave peptide stem crosslinks) (Fig. 3B). The disaccharide-peptide gm-AQK[N]A was detected in high abundance in ORF11 digestion products, confirming that SHEF14 ORF11 possesses dual enzymatic activity and functions both as a glycosyl hydrolase and a D-D endopeptidase, cleaving the bond formed between the D-asparagine residue at the end of the lateral chain and the D-Ala in position 4.

**Fig 3.**
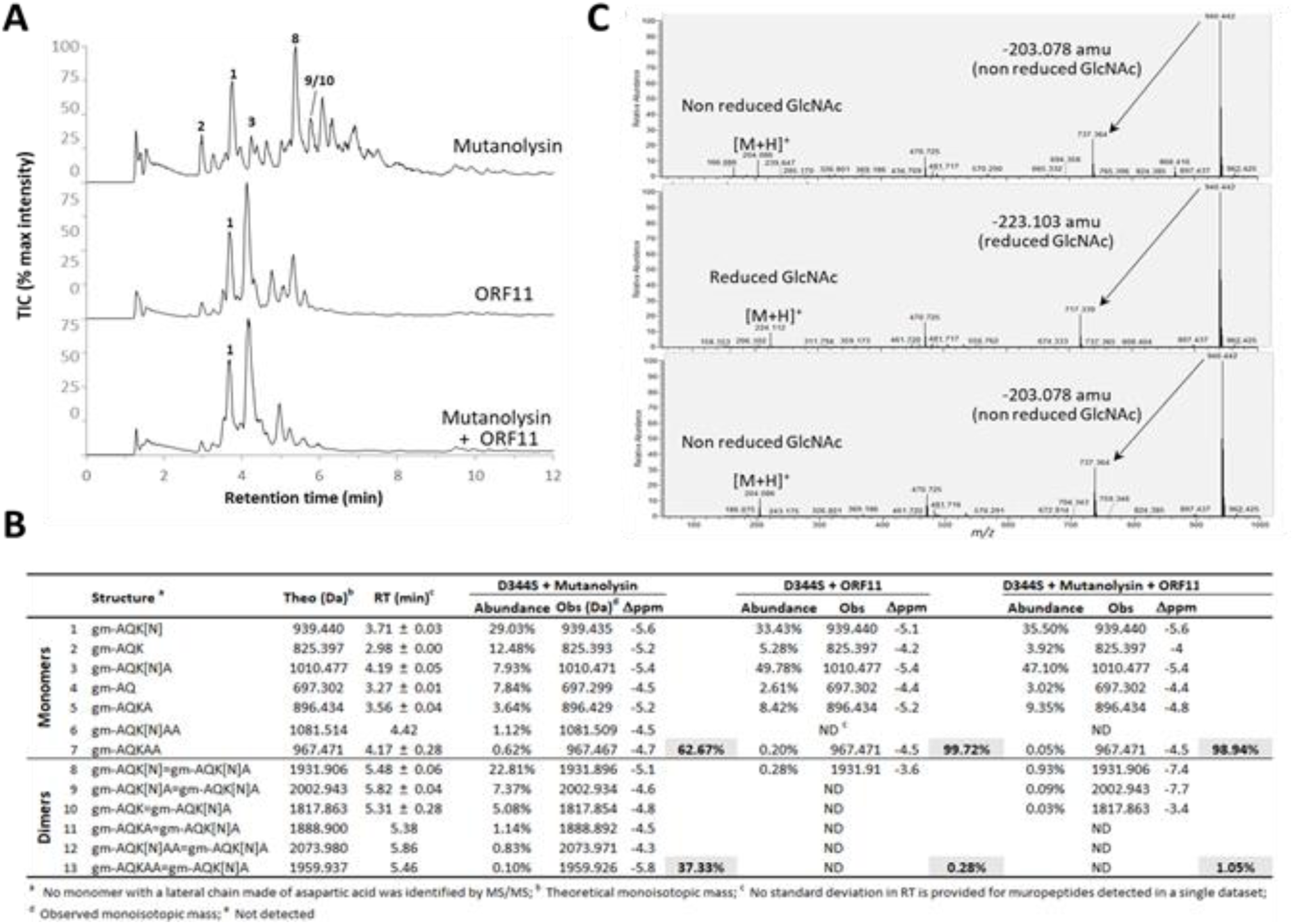
Analysis of ORF11 enzymatic activity using HPLC–MS/MS. (**A**) Total ion chromatograms (TICs) of *E. faecium* D344S cell wall digests treated with mutanolysin, ORF11, or a combination of mutanolysin and ORF11. Major peaks are labelled numerically. (**B**) Relative abundance of muropeptides in each treatment group, corresponding to the peaks indicated in panel A. (**C**) Extracted ion chromatogram of the predominant monomer (peak 1). Loss of a non-reduced GlcNAc residue indicates *N*-acetylmuramidase activity (mutanolysin), whereas loss of a reduced GlcNAc residue indicates *N*-acetylglucosaminidase activity (ORF11).

To define the glycosyl hydrolase specificity of ORF11, we analysed the in-source fragmentation of the major ion corresponding to the gm-AQK[N] monomer (Peak 1) (Fig. 3C) [34]. Unlike mutanolysin-derived ions, which show a 203.078 atomic mass units characteristic of a non-reduced GlcNAc, the peak 1 ion from ORF11 digestion exhibited a 223.103 atomic mass unit loss, which is diagnostic for the removal of reduced GlcNAc. This signature ion demonstrates that ORF11 displays *N*-acetylglucosaminidase activity. Interestingly, whilst ORF11 cleaved the multimeric peptidoglycan fragments, no monosaccharide-peptides (e.g., m-AQK[N] or m-AQK[N]A) were detected, suggesting that the enzyme requires more than a disaccharide motif for activity. This substrate preference likely reflects structural constraints in the D1 active site.

The two predicted catalytic residues in the glucosaminidase and endopeptidase domains (E72A and C523A, respectively) were replaced by alanine residues. Recombinant proteins with single mutations were purified and displayed very low activity against D344S peptidoglycan (data not shown). Their activity was therefore tested on peptidoglycan fragments generated by mutanolysin digestion and results are shown in Fig. S5. As expected, the combined activity of mutanolysin and ORF11 led to a very limited proportion of multimers (26.2% for mutanolysin alone versus 2.3% for mutanolysin + ORF11) due to the endopeptidase activity of the lysin. A similar drastic reduction in dimers was observed in the double digestion with mutanolysin and the E72A ORF11 variant (2.1%), indicating intact endopeptidase activity. The C523A mutation abolished the reduction of the dimer proportion following a double digestion with mutanolysin, showing that domain 4 has endopeptidase activity.

## Discussion

Genes homologous to *orf11* are conserved across *Enterococcus faecium Minhoviruses* and in putative tail proteins from *Staphylococcus* phages such as P68 and Andhra (26% amino acid identity). More distant sequence homology (<20% identity) is also detectable with the PlyCA endolysin from *Streptococcus pyogenes* C1 (Fig. 1D, 2B). The other major human pathogen within the *Enterococcus* genus, *E. faecalis*, is not infected by any known *Minhovirus*; instead, it is targeted by the related podovirus genus *Copernicusvirus*. Although *Copernicusvirus* genomes share ∼50% nucleotide identity overall, the gene equivalent to ORF11 shows <20% amino acid identity to SHEF14 ORF11 (Fig 2B). Notably, *Minhoviruses* along with phage P68 and Andhra, all have ORF11 homologues predicted within the baseplate or the tail spike complex, positioning them for direct interaction with host cell surface structures [25, 28, 30].

Our structural analysis revealed that ORF11 adopts a distinctive four-domain architecture, comprising two catalytic domains displaying glycosyl hydrolase (D1) and peptidase (CHAP, D4) activity closely resembling the corresponding domains in PlyCA. In contrast, the intermediate domains D2 and D3 appear to be specific to *E. faecium Minhoviruses* and are absent from related tail spike proteins in other monoderm podoviruses. Similar dual-domain architectures have been described in tail proteins such as Andhra Gp10, where receptor binding to wall teichoic acids is thought to activate the lytic domains to mediate localized peptidoglycan cleavage [21, 25]. Notably, the ORF11 structural homologue in phage Andhra (Gp10), is a putative dimer in its cryo-EM reconstruction [25].

Since no detailed information is available on the specific bond cleaved by C1_PlyCA or Andhra_Gp10, we performed *in vitro* assays using recombinant ORF11. LC-MS/MS revealed that the enzyme displays both *N*-acetylglucosaminidase and D,D-endopeptidase activities encoded by two separate domains (D1 and D4, respectively). The loss of endopeptidase activity following substitution of the predicted catalytic cysteine with alanine (C523A) underscores the essential role of this residue in peptide-bond cleavage. Although the C523A variant retained measurable glucosaminidase activity, this activity alone did not yield soluble muropeptides or detectable monosaccharide–peptide products. This suggests that the glycosylhydrolase function of ORF11 is intrinsically weak and, in the absence of concurrent endopeptidase activity, unable to release peptidoglycan fragments. Together, these findings support a model in which both catalytic activities of ORF11 act cooperatively, with peptide-bond cleavage likely required to expose or liberate glycan substrates for efficient hydrolysis. Furthermore, the presence of the CHAP-like D3 domain is suggestive of a function in cell wall binding-although our suggests this does not occur in the absence of the other domains. It is also tempting to suggest that the D3 is important in the degradation of *E. faecium* peptidoglycan (D-aspartate-D-asparagine cross bridges) versus that of *E. faecalis* (D-ala-D-ala). However there seems to be more than one evolved solution to cleaving *E. faecium* peptidoglycan since the *Salasmavirus* Athos contains a different type of putative dual-action tail-spike protein (Fig S3).

Despite its clear capacity to hydrolyse purified peptidoglycan, preliminary bactericidal assays showed that ORF11 does not exhibit detectable antimicrobial activity using spot assay on lawns of *E. faecium* E1071, Time-kill curves (Fig. S6 and data not shown). It is possible that the bacteria are protected by the presence of EPA on their surface, precluding access. Another possibility is that the enzyme needs to be part of a larger complex to be active. This result suggests that ORF11 primary role is not in direct cell lysis but rather in the early phases of the phage life cycle, such as local digestion of peptidoglycan to facilitate DNA injection. Based on these observations, we propose that ORF11 functions as an externally functional tail or spike-associated lysin acting specifically at the point of host recognition and entry, similar to that of gp11 from staphylococcus phage p68 (Fig 2B) [23], rather than a canonical endolysin acting at the point of progeny release., acting specifically at the point of host recognition and entry.

EPA is a key determinant of host range in *Enterococcus* phages [13, 16]. Although we found that variations in EPA do not abolish the peptidoglycan-cleaving activity of ORF11, they do influence the host range of phage SHEF14, which is restricted to strain E1071 [13]. This suggests that the substrate specificity of phage SHEF14 and other *Minhoviruses* is not dictated by ORF11 but likely emerges from a more complex interplay between the phage tail machinery and host surface receptors. In this context, ORF11 appears to contribute to cell-wall penetration rather than host recognition, which is probably mediated by additional tail or baseplate components that interact with EPA and define the strain-specific tropism of SHEF14.

Structural homologues for the two central domains of ORF11 (D2 and D3) have not been identified. We initially hypothesized that these domains might participate in interactions with host surface polysaccharides such as EPA or contribute to dimerization, thereby facilitating to correctly position the lytic domains during infection. However, pull-down assays revealed no detectable binding between recombinant D3 (residues N322-Y481) and purified cell walls-although we acknowledge that we do not know the oligomeric state of this construct (Fig. S7). It is also possible that D3 requires either interactions with the rest of the protein domains of Orf11 or cooperative interactions with other tail proteins within the phage structure. Future studies may elucidate how D2 and D3 interface with other components of the *Minhovirus* tail apparatus, with the aim of defining the molecular determinants of host recognition and enabling the rational engineering of phage-based therapeutics targeting multidrug-resistant *E. faecium*.

In conclusion, we report the first crystal structure of a tail-spike protein from an *Enterococcus* phage, ORF11 from the *E. faecium* SHEF14 *Minhovirus*. ORF11 exhibits a distinctive four-domain architecture including two intermediate domains that appear unique to *E. faecium*-infecting *Minhoviruses*. This enzyme displays dual *N*-acetylglucosaminidase and D,D-endopeptidase activities, yet lacks detectable bactericidal activity, consistent with a role as a tail-associated lysin that facilitates early adsorption and local peptidoglycan penetration rather than host cell lysis. Together, these findings provide new insight into *Minhovirus*–host interactions and offer a structural and biochemical foundation for developing phage-based strategies to combat multidrug-resistant *E. faecium*.

## Materials and Methods

### Bacterial strains, plasmids, oligonucleotides and growth conditions

Bacterial strains, plasmids and oligonucleotides are described in Table S1. *E. faecium* strains were grown on THY (Todd Hewitt supplemented with 1% [w/v] yeast extract) agar plates or in THY broth. *E. coli* was grown on LB-agar plates or in LB broth at 37°C. When needed, ampicillin (100 µg/mL) or chloramphenicol (35 µg/mL) was added.

### Construction of plasmids for protein expression

The DNA fragment encoding ORF11 was amplified with oligonucleotides ORF11_Fw and ORF11_Rev using Phusion polymerase (Fisher scientific) and phage 14 DNA as a template. The PCR product was cloned into pET21a(+) via the NdeI and XhoI sites. A synthetic DNA fragment flanked by NcoI and SacII restriction sites was cloned into pET21_S to produce recombinant ORF11 domain 3 (residues N322-Y481) with a C-terminal Strep-tag. The synthetic DNA encoded a fusion between the predicted signal peptide of *R. Johnstonii* 3841 gene *RL1724* (30 residues) fused to the sequence encoding ORF11 residues. The resulting sequence is described in Fig. S8.

### Protein expression and purification

Recombinant proteins were expressed in *E. coli* BL21(DE3) grown in auto induction media or M9 for selenomethionine labelling [35]. Cells were inoculated at OD600=0.025 and grown at 37°C until the OD_600_ reached 0.5. Cultures were then cooled down and incubated overnight at 20°C, after addition of 0.5mM IPTG.

For purification, cells were harvested by centrifugation, resuspended in a lysis buffer (50 mM Tris-HCl pH 8, 300 mM NaCl) and disrupted twice using a French Press (1200 psi). Following centrifugation (45,000 x g, 20 min, 4°C), the soluble fraction was applied onto a 5 mL Ni-Nta (ORF11) or a StrepTrap XT (domain 3) column using an Äkta Pure system (Cytiva). The resin was washed with 50 mM Tris-HCl pH 8, 300 mM NaCl and the proteins were eluted in a single step using Tris-HCl pH 8, 300 mM NaCl, and 500 mM imidazole (ORF11) or Tris-HCl pH 8, 300 mM NaCl, and 2.5 mM desthiobiotin (domain 3). The fractions containing the protein of interest were pooled and further purified on a Superdex 200 16/600 column (Cytiva). Pure fractions, as analyzed by SDS-PAGE, were pooled.

### Bioinformatic analysis

Genome regions were compared using clinker: https://cagecat.bioinformatics.nl/tools/clinker [36]. Structural predictions were enabled using the AlphaFold server [37] and structural homologues via FoldSeek [38]. Structural comparisons and images were generated with ChimeraX [39].

### Crystallization, data-collection and analysis

Following initial vapour diffusion crystallization trials employing commercial screens on a Mosquito liquid handling robot (SPT labtech), crystals with a plate-like morphology were grown using purified ORF11 at 10 mg/mL and 17% (w/v) PEG3350, 0.1M MES buffer pH 6.7. Crystal size was improved using seeding. The crystals were sent to the Diamond Light Source at Harwell near Oxford for data collection on beamlines i03 and i04. Processing was carried out with DIALS, and initial molecular replacement attempts were performed using PHASER [40] from the CCP4 [41] suite with models produced using AlphaFold for full size ORF11 and models of individual domains. These attempts proved unsuccessful, and phases were ultimately determined by SAD from the selenium anomalous signal of selenomethionine incorporated protein crystals from data collected at 0.9795 Å wavelength (Table 1). The substructure and initial protein model were determined from the xia2.multiplex processed data using Crank2 to create a preliminary protein model of ∼90% completeness. Subsequent model building and refinement were carried out using iterations of COOT [42] and Isolde [43], then Refmac5 from the CCP4 suite and validated using MolProbity [44].

**Table 1.** Data collection for ORF11 (SeMet data).

|  |  |
| --- | --- |
| Space group | P2<sub>1</sub>2<sub>1</sub>2<sub>1</sub> |
| Unit cell Å ; ° | a= 72.8 b= 88.0 c= 233.8; α=β=γ= 90 |
| Resolution Å | 69.51-2.50 (2.58-2.50) |
| Rmerge (all I<sup>+</sup> & I<sup>-</sup>) | 0.113 (0.942) |
| Rpim (all I<sup>+</sup> & I<sup>-</sup>) | 0.032 (0.264) |
| Mean ((I) / sd (I)) | 17.1 (2.4) |
| Completeness % | 100 (100) |
| Multiplicity | 13.5 (13.6) |
| CC half | 0.89 (0.77) |
| Anomalous completeness % | 100 (100) |
| Anomalous multiplicity | 7.1 (7.0) |
| Total observations | 716614 (61802) |
| Total unique | 52998 (4533) |
| R-factor (%) | 0.217 |
| R<sub>free</sub> (%) | 0.24 |
| RMSD - bonds | 0.0031 |
| - angles | 1.154 |
| No. refined atoms (protein/solvent) | 10430 (10153/277) |
| PDB entry code | 9T7R |

### Peptidoglycan extraction

*E. faecium* strains were grown overnight in 900 mL of THY broth from a single colony. Cells were spun and the cell pellet was snap frozen in liquid nitrogen. Cell walls were extracted by resuspending the cells in 20 mL of boiling MilliQ water (MQ) before the addition of 5 mL warm 20% (w/v) SDS (4% final concentration). After 30 min at 100°C, the cells were allowed to cool down to room temperature. Insoluble cell walls were recovered at 20,000 x *g* for 10 min and washed five times using warm MQ water. Proteins covalently bound to peptidoglycan were removed by pronase E treatment (final concentration of 2 mg/mL, 4 hrs at 60°C). Protease-treated cell walls were washed 6 times with 30 mL of MQ water before covalently bound polymers were removed by incubation in 1 M HCl for 5 hrs at 37°C. Insoluble pure peptidoglycan was washed 6 times with MQ water, snap frozen in liquid nitrogen, freeze-dried and resuspended at a final concentration of 10 mg/mL.

### Peptidoglycan digestion and preparation of soluble muropeptides

1mg of peptidoglycan was digested with 100U of mutanolysin (Merck) in 10 mM phosphate buffer (pH 5.5) or 100 µg of ORF11 in phosphate buffer saline (10 mM Na_2_PO_4_, 3mM KCl and 150 mM NaCl, pH 7.4) in a final volume of 100 µL. After 16h incubation at 37°C, enzymes were inactivated 5 min at 100°C. Soluble fragments were recovered after centrifugation at 25,000 x *g* for 10 min. Disaccharide-peptides were reduced by adding one volume of 200 mM borate buffer (pH 9.0) and 500 µg of sodium borohydride. After 20 min at room temperature, the pH was adjusted to 4.5 using phosphoric acid.

### LC-MS/MS

LC-MS analysis was carried out using an Ultimate 3000 Ultra High-Performance Chromatography system (UHPLC; Dionex/Thermo Fisher Scientific) coupled with a Orbitrap Exploris 240 (Thermo Fisher Scientific). Muropeptides were separated using a C18 analytical column (Hypersil Gold aQ, 1.9 µm particles, 150 × 2.1 mm; Thermo Fisher Scientific) at a temperature of 50°C. Elution was performed at 0.25 mL/min by applying a mixture of solvent A (water, 0.1% [v/v] formic acid) and solvent B (acetonitrile, 0.1% [v/v] formic acid). LC conditions were 0–12.5% B for 25 min increasing to 20% B for 10 min. After 5 min at 95%, the column was re-equilibrated for 10 min with 100% buffer A.

The Orbitrap Exploris 240 was operated under electrospray ionization (H-ESI high flow)-positive mode, full scan (m/z 150–2250) at resolution 120,000 (FWHM) at m/z 200, with normalized AGC Target 100%, and automated maximum ion injection time (IT). Data-dependent MS/MS were acquired on a ‘Top 5’ data-dependent mode using the following parameters: resolution 30,000; AGC 100%, automated IT, with normalized collision energy 25%. Raw data analysed in Fig. 3 are available through the GlycoPost repository (Project reference GPST000649).

### Peptidoglycan LC-MS data analysis

Raw data was analysed using the proprietary software Byos® (Protein Metrics) to identify monomers based on MS/MS data. The fasta file and search parameters used are described in Fig. S9A and S9B. The disaccharide-peptides identified with Byos® were used to build the database described in Fig. S9C. Muropeptides were identified in deconvoluted datasets with PGFinder v1.2.0 using default parameters.

### Antimicrobial assays

Exponential-phase *E. faecium* E1071 was diluted to 5 10^5^ CFU/mL, and 100 µL of cells were incubated with recombinant ORF11 at 2-fold serial dilutions ranging from 256 µg/mL to 0.25 µg/mL. A sample without ORF11 served as the negative control. Aliquots of the mixtures were taken at different time points (0 h, 1 h, and 3 h) and 1 µL (equivalent to 50 CFUs) was placed onto THY agar plates and incubated overnight.

### Pull-down assay

Various amounts of *E. faecium* E1071 cell wall (0, 10, 30, 100, 300, 1000, and 3000 µg) were incubated in the presence of 20 µg of purified domain D3 or LysM (used as a positive control) in 50 µL PBS. After 20 min at room temperature, the mixtures were centrifuged at 13,300 rpm for 6 min, and 30 µL of the supernatant was mixed with 7.5 µL of 5x SDS-PAGE loading buffer containing 100 mM DTT. Samples were heated at 100°C for 10 min, and 10 µL were then loaded onto a 12% SDS-PAGE gel.

## Data Availability Statement

Structural co-ordinates are deposited in the PDB under accession 9T7R and will be released on publication. All other data is available on request. Code for PGFinder code is available via https://github.com/Mesnage-Org/pgfinder.

## Acknowledgements

SM was funded by a BBSRC grant (BB/w013800/1). JW was funded by a BBSRC fellowship (BB/Z515188/1), ES was funded by a NIHR Sheffield Biomedical Research Centre (BRC) studentship (NIHR203321). AA was supported via funding from Prince Sattam bin Abdulaziz University project number (PSAU/2026/R/1447). CE was supported by EPSRC grant EP/E036252/1. YZ and JL were supported by a grant from the National Natural Science Foundation of China (32322082).

## Author Contributions

**(CreDIT attributions): SM**: Conceptualisation, Data curation, Funding acquisition, Investigation, Methodology, Supervision, Validation, Visualization, Writing- original draft, review and editing

**JSW:** Investigation, methodology, Writing- review and editing

**YZ**: Methodology, Investigation, Visualization, Writing- review and editing

**AA**: Methodology, Writing- review and editing

**MR**: Investigation, methodology, Writing- review and editing

**ES**: Investigation, Writing- review and editing

**CE**: Investigation, Methodology, Writing- review and editing

**JL**: Funding acquisition, Writing- review, supervision, Writing- review and editing

**JBR**: Conceptualisation, Data curation, Investigation, Methodology, Supervision, Visualization, Writing- original draft, review and editing

**GPS**: Conceptualisation, Data curation, Funding acquisition, Investigation, Methodology, Supervision, Validation, Visualization, Writing- original draft, review and editing

---

**Fig. S1.**
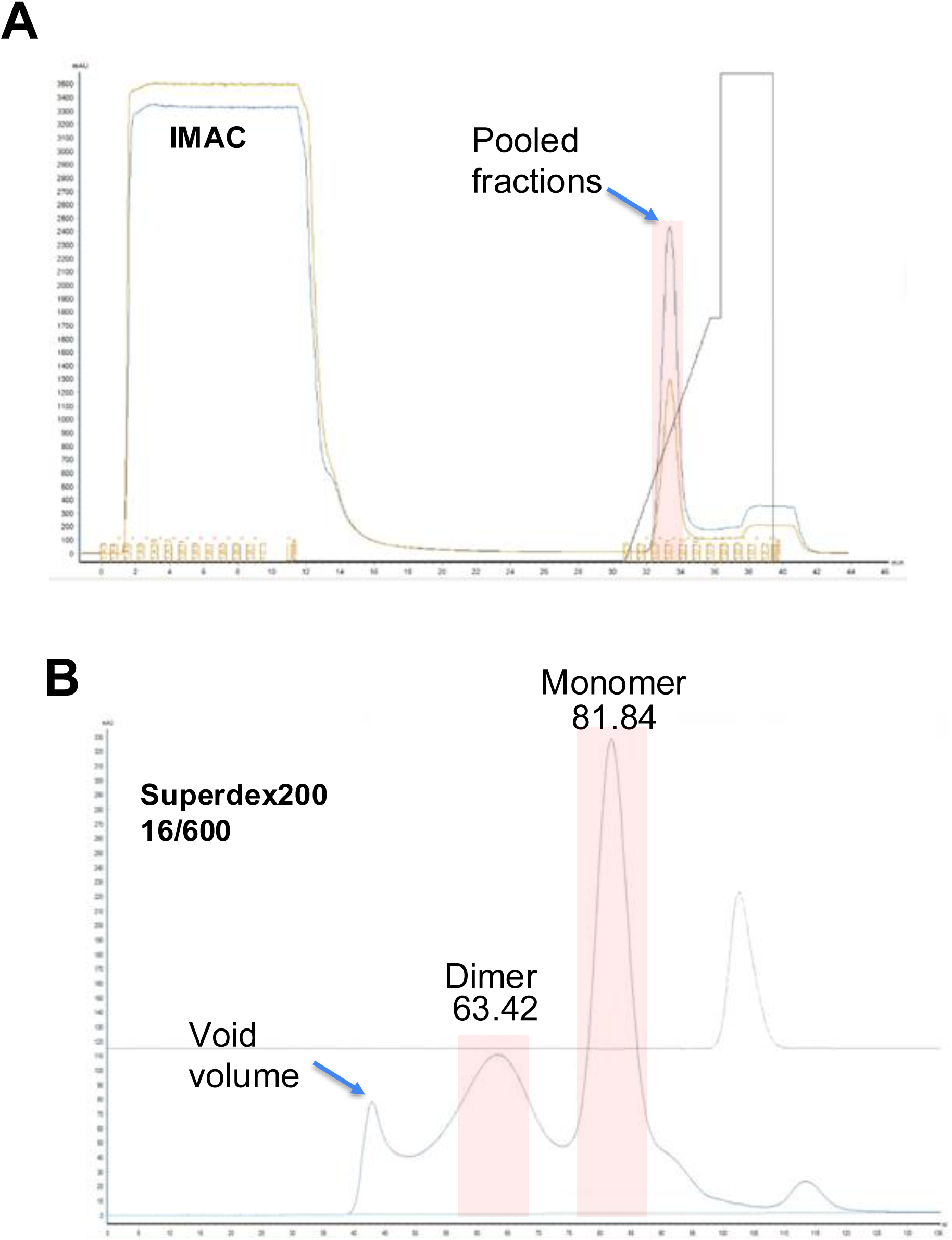
Purification of ORF11. **A**, Immobilized Metal Affinity Chromatography (IMAC) shows a single peak of proteins eluted with approximately 150mM imidazole. **B**, Gel filtration purification of the fractions pooled (red area, ∼15mg of proteins in 3.5mL) were injected in a volume of 3.5mL on a Superdex200 16/600 gel filtration column. Retention times of protein standards (IgG[ 158kDa] at 64.47min and human albumin [66kDa] at 74.45 min) suggest that the two peaks observed for ORF11 correspond to a dimer and monomer form of the protein.

**Fig. S2.**
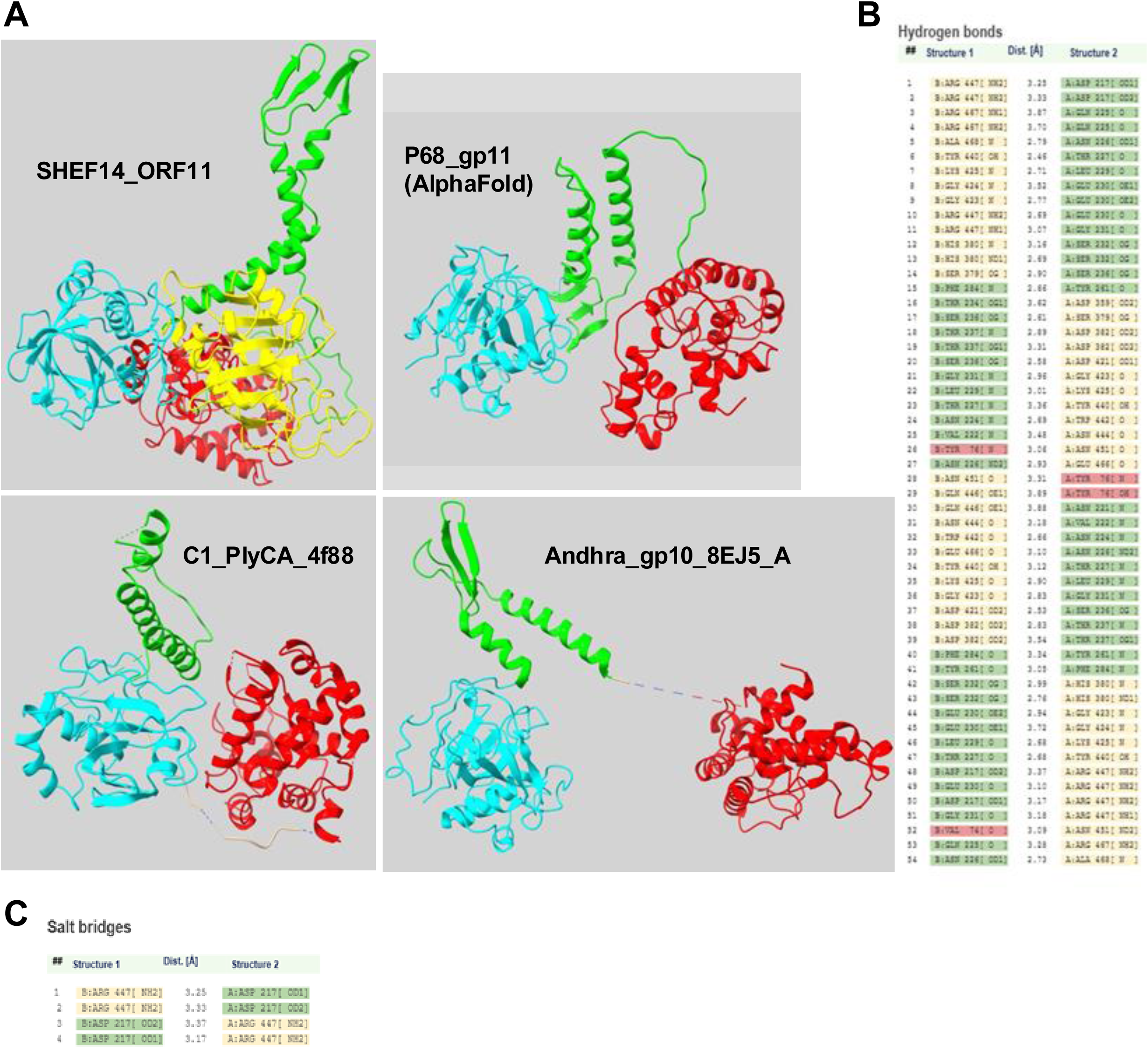
Comparison of SHEF14_ORF11 with structural homologues and analysis of dimer interface. **A**, structure of ORF11 and homologs. GH domains in red, CHAP in cyan and the linker domain in lime. The D3 domain of ORF11 is in yellow. Regions with no structure in the PDB are dashed. **B/C,** Analysis of the ORF11 dimer interface: PISA server analysis of interactions formed within ORF11 dimer interface (D2-D3). (A) Salt bridges and (B) hydrogen bonds between chain A and B are shown, and coloured based on the domain architecture shown in Figure 1c.

**Fig. S3.**
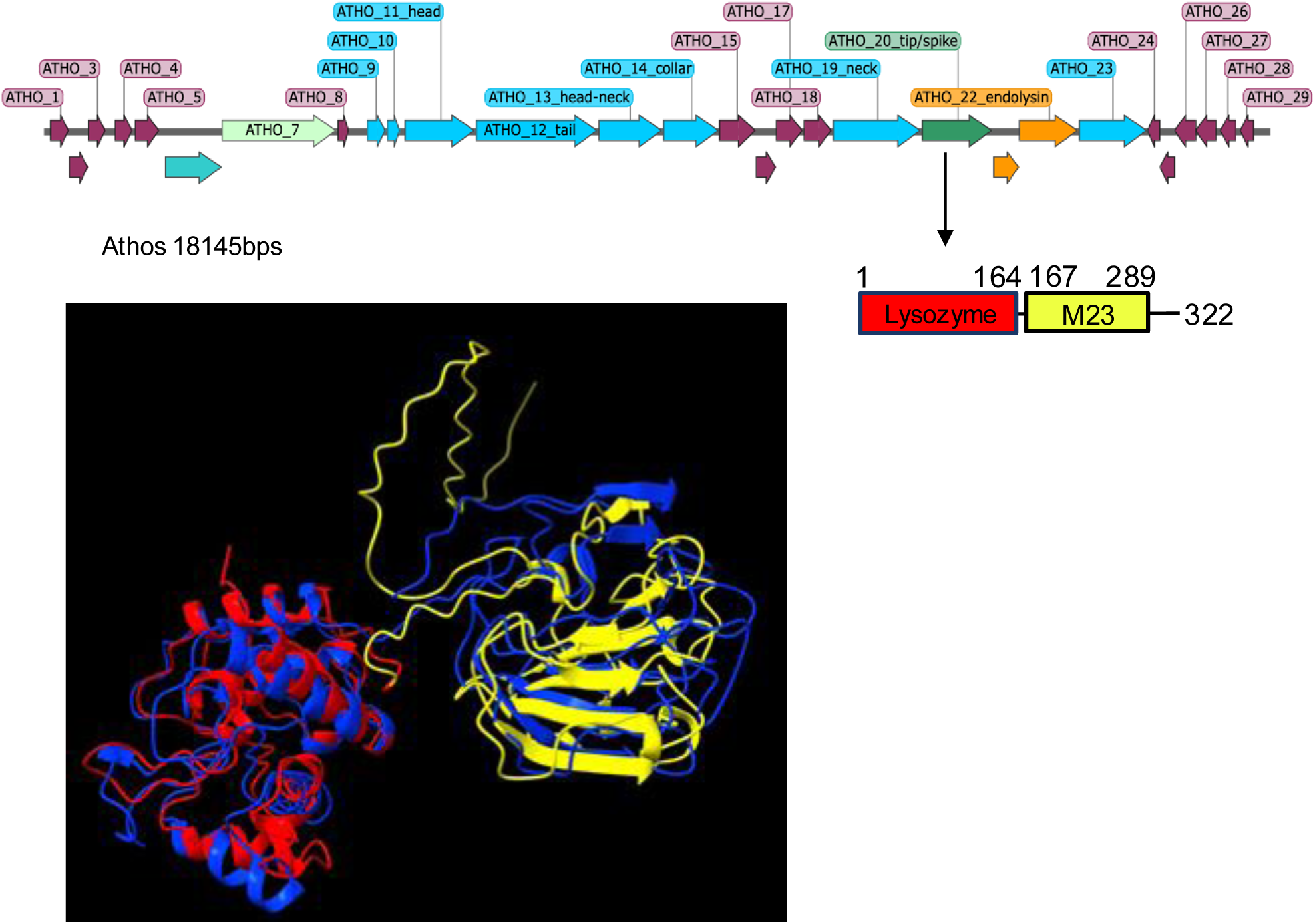
Dual domain spike hydrolase encoded by the *Salasmavirus* Athos. Schematic of Phage_Athos genome with orf20 in green (top), indicating location of Lysozyme and M23 peptidase domains as rectangles; with a structural alignment (lower left) of Athos_20 (red/ yellow domains) with 3SCQ-phi29_gp13 tail tip (blue)

**Fig. S4.**
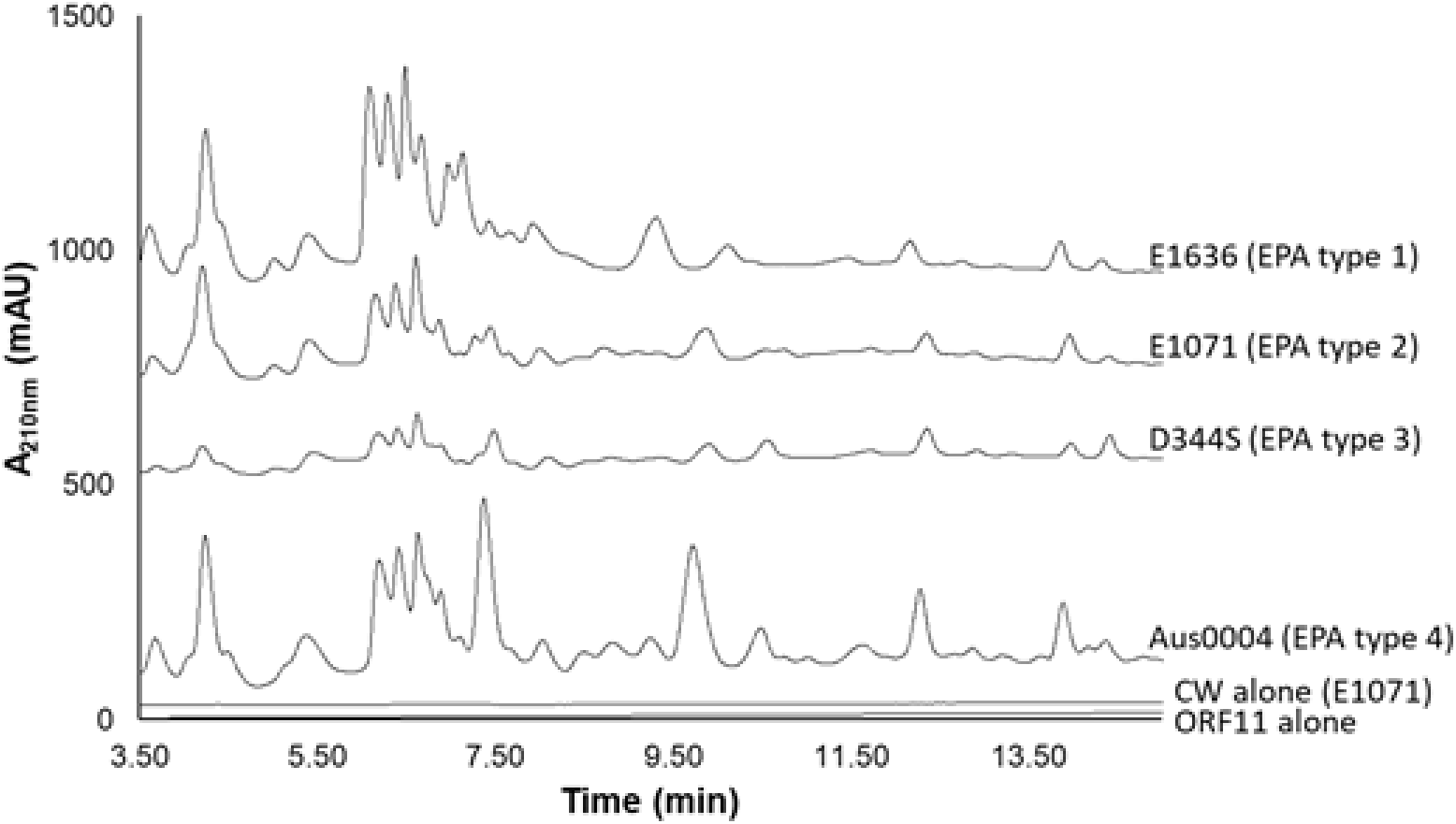
HPLC muropeptide profiles of *E. faecium* EPA variant cell walls digested with ORF11. Cell walls were extracted from four representative EPA variant strains and 500 µg were digested with 50 µg of recombinant ORF11 for 16h at 37°C in a final volume of 100 µL. Soluble muropeptides equivalent to 25 µL of the digestion products were separated by rp-HPLC and detected by monitoring UV absorbance at 210nm. Cell walls from strains E1636 (EPA variant 1), E1071 (EPA variant 2), D344S (EPA variant 3), and Aus0004 (EPA variant 4) were used as substrates.

**Fig. S5.**
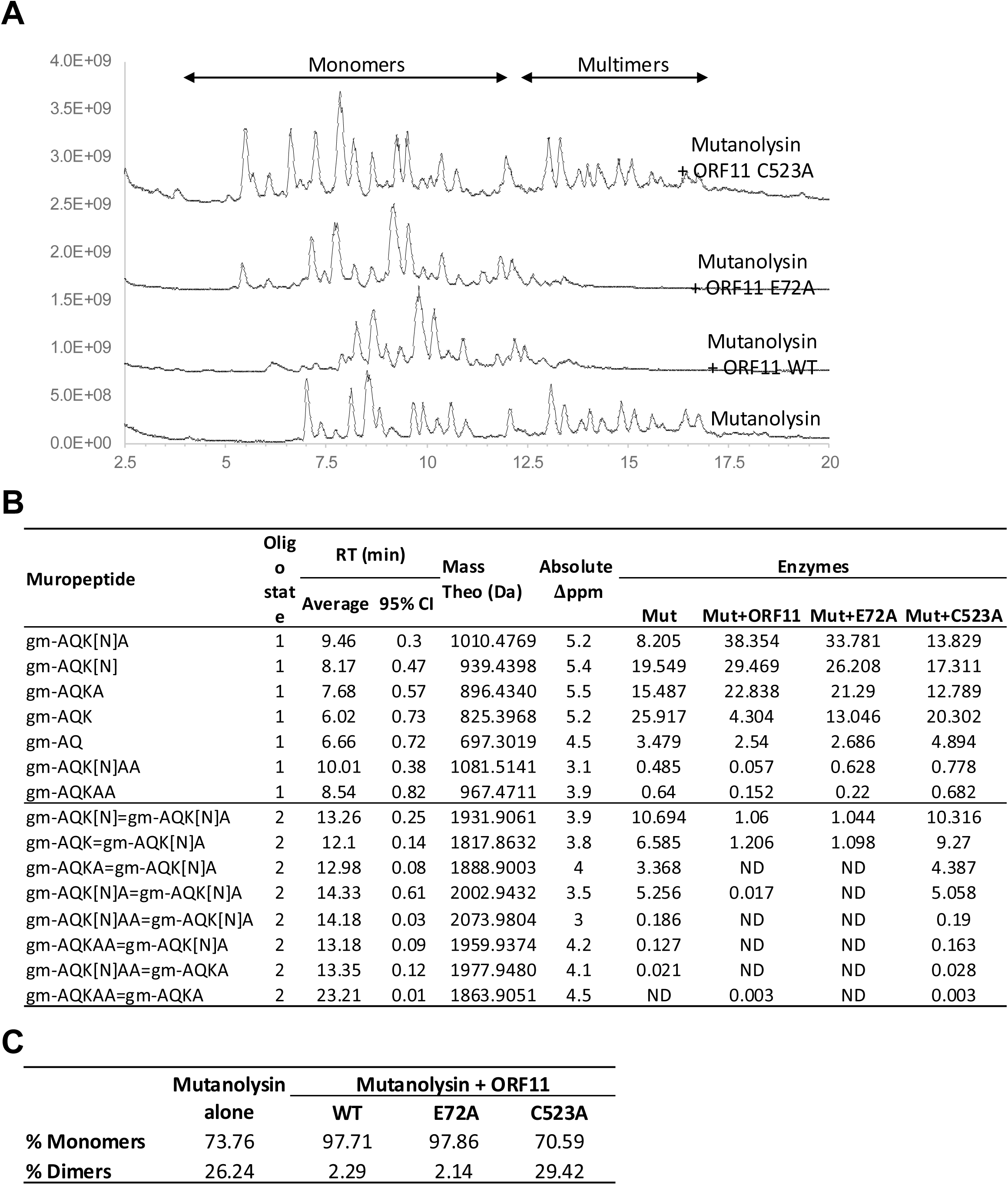
HPLC muropeptide profiles of *E. faecium* E1071 cell walls digested with mutanolysin and ORF11. 500 µg of cell walls were digested with 50U of Mutanolysin and 125µG of purified ORF11 WT, E72A or C523A mutant. for 16h at 37°C in a final volume of 100 µL. Soluble muropeptides equivalent to 25 µL of the digestion products were analyed by LC-MS/MS. (**A**) Total ion traces corresponding to each digestion. (**B**) PGFinder output of the muropeptide search using the same strategy as in Fig. 3. (**C**) Summary table describing the percentage of monomers and dimers identified by PGFinder.

**Fig. S6.**
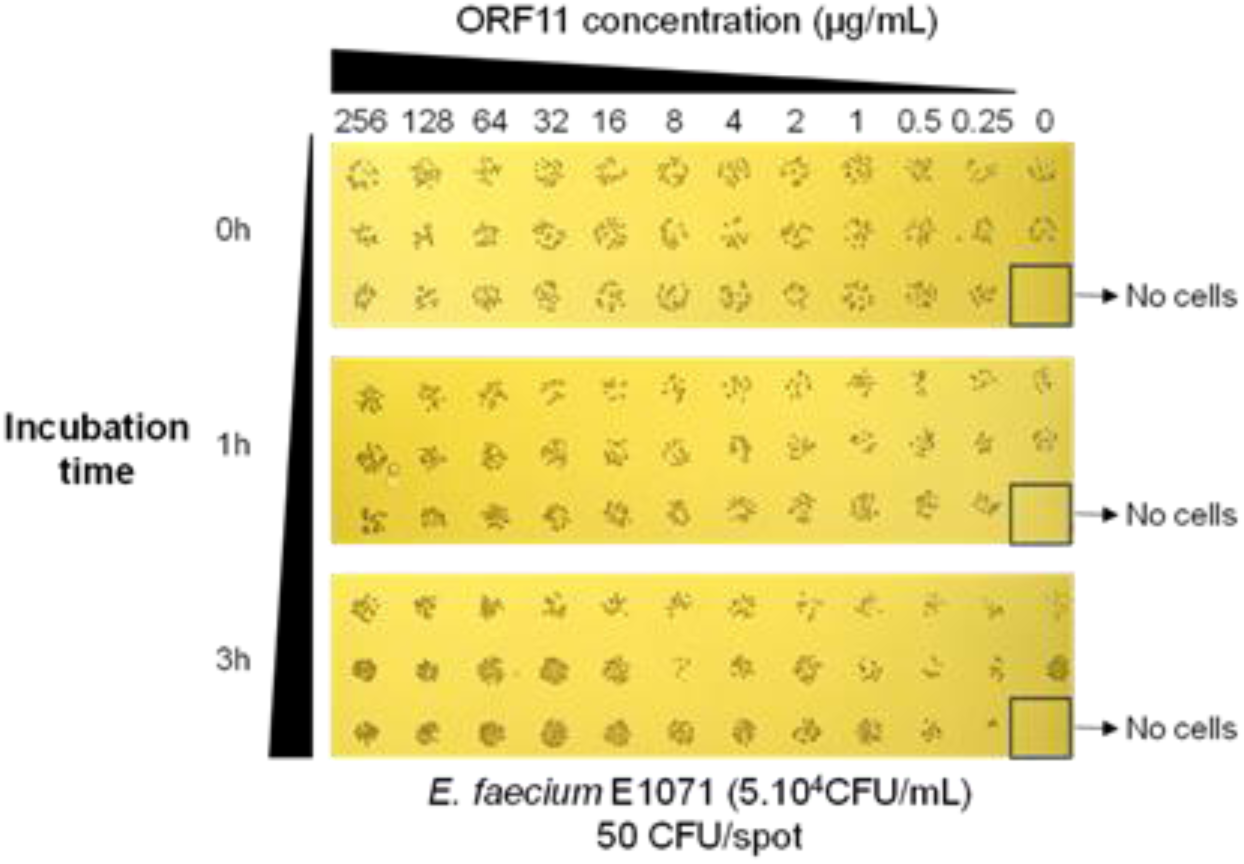
Antimicrobial assay with recombinant ORF11. The equivalent of 5.10^4^ CFUs in 100 µL were incubated in the presence of various concentrations of recombinant ORF11 (from 256 µg/mL to 0.25 µg/mL) for various amounts of time (0, 1 or 3h) and one µL of the mixture was stamped on THY-agar.

**Fig. S7.**
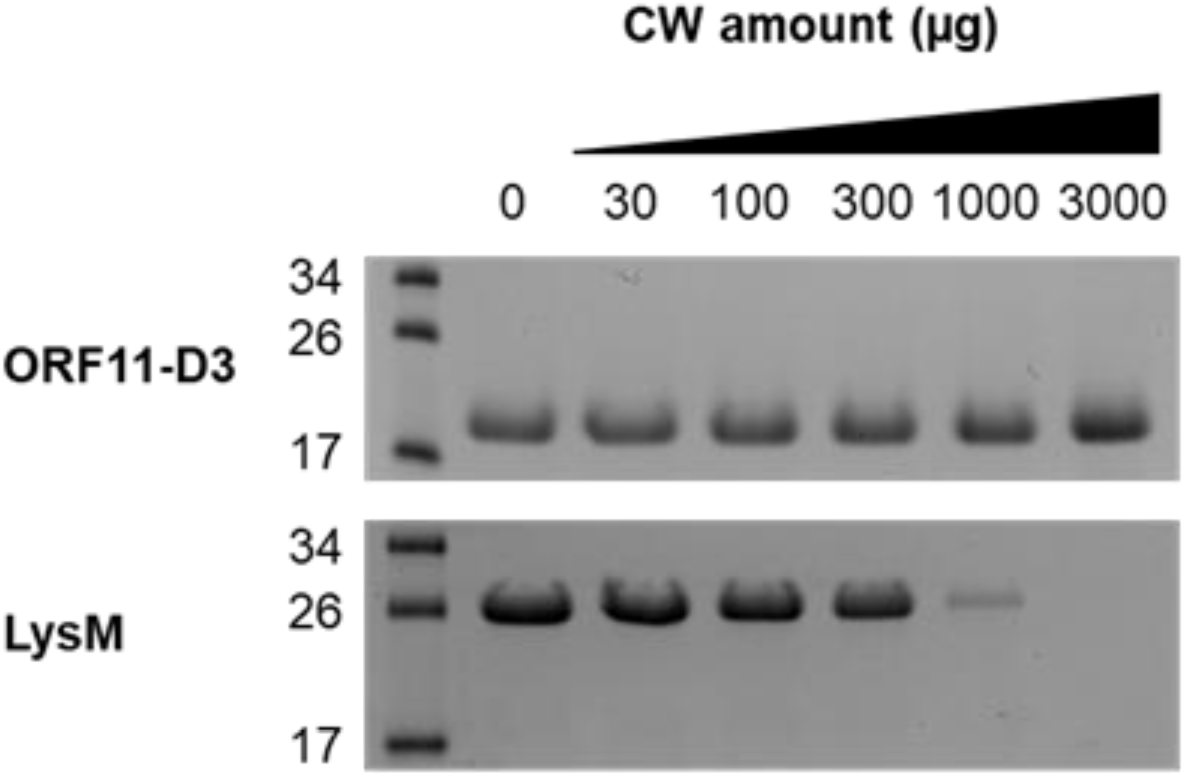
Cell wall binding assay using ORF11 domain 3 and LysM. Various amounts of cell walls purified from *E. faecium* E1071 were incubated in the presence of recombinant ORF11 domain 3 (ORF11-D3) or LysM as a positive control. After an incubation of 20 min at room temperature, cell walls were sedimented and unbound proteins were loaded on a SDS-PAGE. The lack of protein on the Coomassie-stained gel indicates binding to cell walls.

**Fig. S8.**
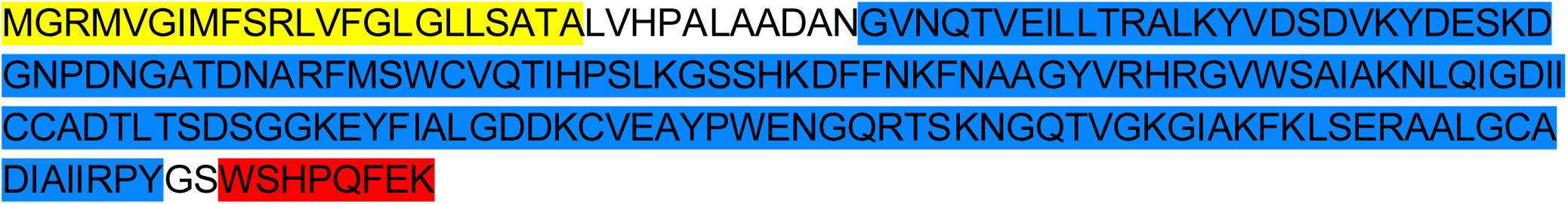
Amino acid sequence of ORF11 domain 3 expressed in *E. coli*. The signal peptide of *R. johnstonii*3841 RL1724 (yellow) followed by 3 residues (ADA) was fused to ORF11 domain 3 (blue; residues N322-Y481). The corresponding polypeptide is fused to a C-terminal Strep-tag (red).

**Fig. S9.**
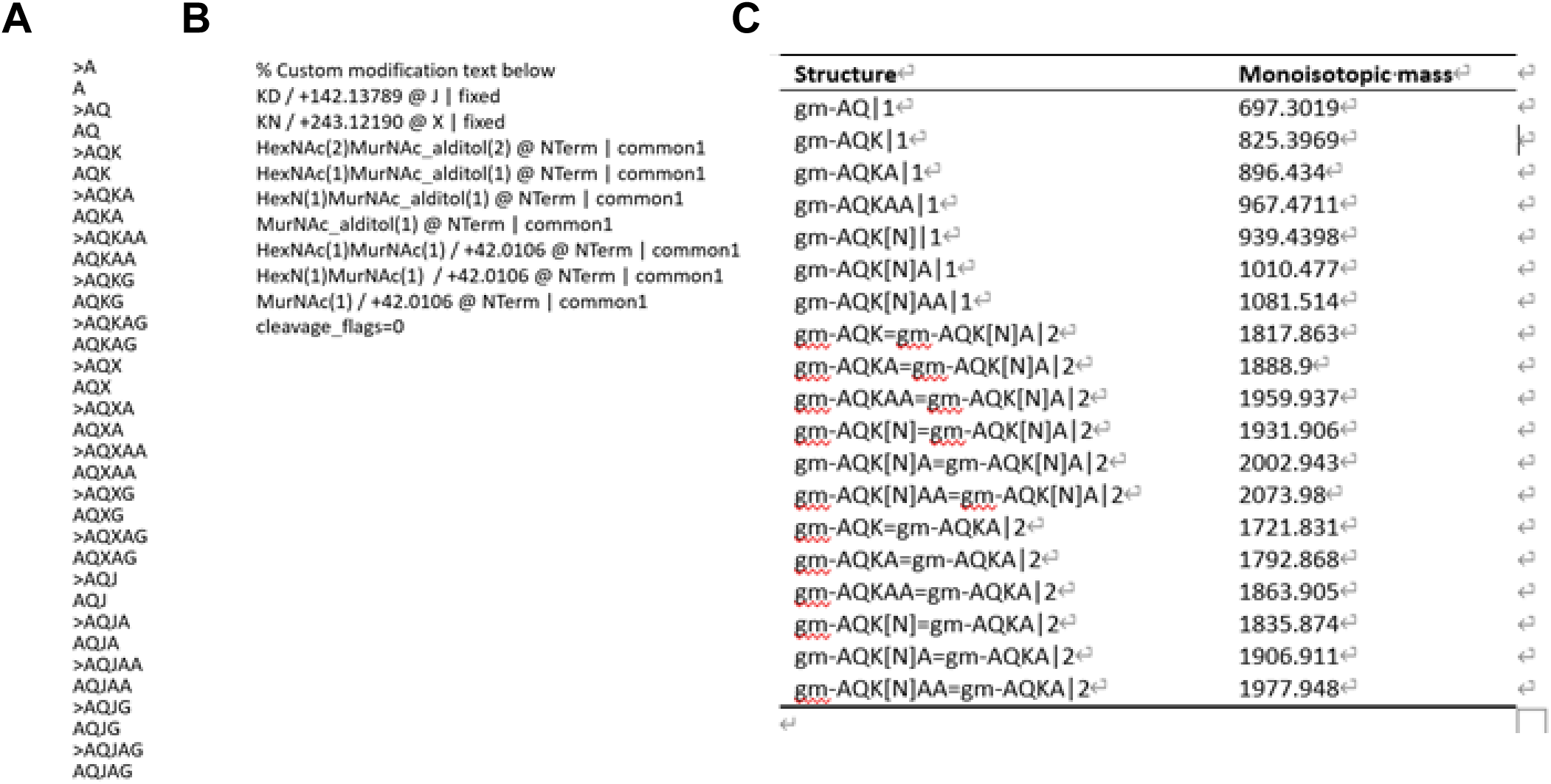
Search parameter for LC-MS/MS analysis using Byos^®^. (A) Fasta file description; J corresponds to a lysine substituted with an aspartate residue and X corresponds to a lysine substituted with an asparagine residue. (B) Description of custom modifications for sugar moieties. (C) Muropeptide list of muropeptides in the database DB_Fm containing monomers and dimers with their corresponding monoisotopic masses.

**Table S1.**
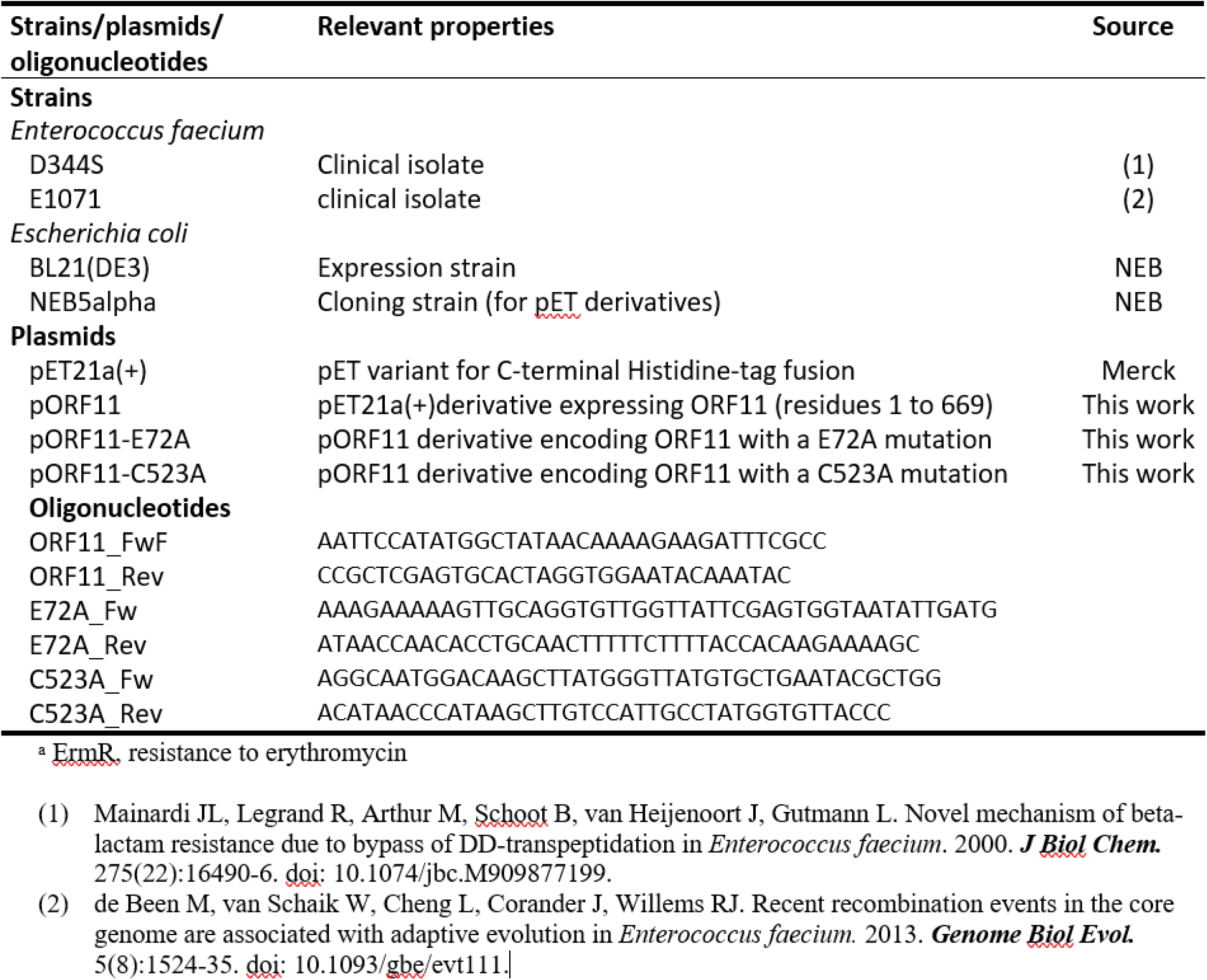
Bacterial strains. Plasmids and oligonucleotides.

## References

1 García-Solache, M. and Rice, L. B. (2019) The Enterococcus: a Model of Adaptability to Its Environment. Clin Microbiol Rev 32, e00058–18 10.1128/CMR.00058-18

2 Guzman Prieto, A. M., Van Schaik, W., Rogers, M. R. C., Coque, T. M., Baquero, F., Corander, J., et al. (2016) Global Emergence and Dissemination of Enterococci as Nosocomial Pathogens: Attack of the Clones? Front. Microbiol. 7 10.3389/fmicb.2016.00788

3 Fiore, E., Van Tyne, D. and Gilmore, M. S. (2019) Pathogenicity of Enterococci. Microbiol Spectr (Fischetti, V. A., Novick, R. P., Ferretti, J. J., Portnoy, D. A., Braunstein, M., and Rood, J. I., eds.) 7, 7.4.9 10.1128/microbiolspec.GPP3-0053-2018

4 Krawczyk, B., Wityk, P., Gałęcka, M. and Michalik, M. (2021) The Many Faces of Enterococcus spp.—Commensal, Probiotic and Opportunistic Pathogen. Microorganisms 9, 1900 10.3390/microorganisms9091900

5 Chatterjee, A., Johnson, C. N., Luong, P., Hullahalli, K., McBride, S. W., Schubert, A. M., et al. (2019) Bacteriophage Resistance Alters Antibiotic-Mediated Intestinal Expansion of Enterococci. *Infect Immun* (Whiteley, M., ed.) 87, e00085–19 10.1128/IAI.00085-19

6 Leclercq, R., Derlot, E., Duval, J. and Courvalin, P. (1988) Plasmid-Mediated Resistance to Vancomycin and Teicoplanin in Enterococcus Faecium. N Engl J Med 319, 157–161 10.1056/NEJM198807213190307

7 Hautemanière, A., Hunter, P. R., Diguio, N., Albuisson, E. and Hartemann, P. (2009) A prospective study of the impact of colonization following hospital admission by glycopeptide-resistant Enterococci on mortality during a hospital outbreak. American Journal of Infection Control 37, 746–752 10.1016/j.ajic.2009.02.007

8 Diallo, K. and Dublanchet, A. (2022) Benefits of Combined Phage-Antibiotic Therapy for the Control of Antibiotic-Resistant Bacteria: A Literature Review. Antibiotics (Basel, Switzerland) 11 10.3390/antibiotics11070839

9 Al-Zubidi, M., Widziolek, M., Court, E. K., Gains, A. F., Smith, R. E., Ansbro, K., et al. (2019) Identification of Novel Bacteriophages with Therapeutic Potential That Target Enterococcus faecalis. *Infection and Immunity* (Whiteley, M., ed.) 87 10.1128/IAI.00512-19

10 Wandro, S., Oliver, A., Gallagher, T., Weihe, C., England, W., Martiny, J. B. H., et al. (2018) Predictable Molecular Adaptation of Coevolving Enterococcus faecium and Lytic Phage EfV12-phi1. Frontiers in microbiology 9, 3192 10.3389/fmicb.2018.03192

11 Mangalea, M. R. and Duerkop, B. A. (2020) Fitness trade-offs resulting from bacteriophage resistance potentiate synergistic antibacterial strategies. Infection and immunity **online ear** 10.1128/IAI.00926-19

12 Lossouarn, J., Beurrier, E., Bouteau, A., Moncaut, E., Sir Silmane, M., Portalier, H., et al. (2024) The virtue of training: extending phage host spectra against vancomycin-resistant *Enterococcus faecium* strains using the Appelmans method. *Antimicrob Agents Chemother* (Tamma, P. D., ed.) 68, e01439–23 10.1128/aac.01439-23

13 Alrafaie, A. M., Pyrzanowska, K., Smith, E. M., Partridge, D. G., Rafferty, J., Mesnage, S., et al. (2024) A diverse set of Enterococcus-infecting phage provides insight into phage host-range determinants. Virus Res 347, 199426 10.1016/j.virusres.2024.199426

14 Evans, E., Ojima, S., Yamashita, W., Kiga, K. and Van Tyne, D. (2026) Characterization of *Enterococcus faecium* -Targeting *Minhovirus* Bacteriophages. *PHAGE: Therapy*, Applications, and Research 26416549251413133 10.1177/26416549251413133

15 Wandro, S., Ghatbale, P., Attai, H., Hendrickson, C., Samillano, C., Suh, J., et al. (2022) Phage Cocktails Constrain the Growth of *Enterococcus*. *mSystems* (Gaglia, M. M., ed.) 7, e00019–22 10.1128/msystems.00019-22

16 Davis, J. L., Norwood, J. S., Smith, R. E., O’Dea, F., Chellappa, K., Rowe, M. L., et al. (2025) Dissecting the Enterococcal Polysaccharide Antigen (EPA) structure to explore innate immune evasion and phage specificity. Carbohydrate Polymers 347, 122686 10.1016/j.carbpol.2024.122686

17 De Been, M., Van Schaik, W., Cheng, L., Corander, J. and Willems, R. J. (2013) Recent Recombination Events in the Core Genome Are Associated with Adaptive Evolution in Enterococcus faecium. Genome Biology and Evolution 5, 1524–1535 10.1093/gbe/evt111

18 Zinder, N. D. and Lederberg, J. (1952) GENETIC EXCHANGE IN SALMONELLA. J Bacteriol 64, 679–699 10.1128/jb.64.5.679-699.1952

19 Demerec, M. and Fano, U. (1945) BACTERIOPHAGE-RESISTANT MUTANTS IN ESCHERICHIA COLI. Genetics 30, 119–136 10.1093/genetics/30.2.119

20 Nelson, D., Schuch, R., Zhu, S., Tscherne, D. M. and Fischetti, V. A. (2003) Genomic Sequence of C_1_ , the First Streptococcal Phage. J Bacteriol 185, 3325–3332 10.1128/JB.185.11.3325-3332.2003

21 Cater, K., Dandu, V. S., Bari, S. M. N., Lackey, K., Everett, G. F. K. and Hatoum-Aslan, A. (2017) A Novel *Staphylococcus* Podophage Encodes a Unique Lysin with Unusual Modular Design. mSphere (Fey, P. D., ed.) 2, e00040–17 10.1128/mSphere.00040-17

22 Alrafaie, A. M. and Stafford, G. P. (2023) Enterococcal bacteriophage: A survey of the tail associated lysin landscape. Virus research 327, 199073 10.1016/j.virusres.2023.199073

23 Hrebík, D., Štveráková, D., Škubník, K., Füzik, T., Pantůček, R. and Plevka, P. (2019) Structure and genome ejection mechanism of Staphylococcus aureus phage P68. Sci Adv 5, eaaw7414 10.1126/sciadv.aaw7414

24 Xu, J., Wang, D., Gui, M. and Xiang, Y. (2019) Structural assembly of the tailed bacteriophage ϕ29. Nat Commun 10, 2366 10.1038/s41467-019-10272-3

25 Hawkins, N. C., Kizziah, J. L., Hatoum-Aslan, A. and Dokland, T. (2022) Structure and host specificity of *Staphylococcus epidermidis* bacteriophage Andhra. Sci. Adv. 8, eade0459 10.1126/sciadv.ade0459

26 Lavigne, R., Seto, D., Mahadevan, P., Ackermann, H.-W. and Kropinski, A. M. (2008) Unifying classical and molecular taxonomic classification: analysis of the Podoviridae using BLASTP-based tools. Research in Microbiology 159, 406–414 10.1016/j.resmic.2008.03.005

27 Chen, W., Xiao, H., Wang, L., Wang, X., Tan, Z., Han, Z., et al. (2021) Structural changes in bacteriophage T7 upon receptor-induced genome ejection. Proc. Natl. Acad. Sci. U.S.A. 118, e2102003118 10.1073/pnas.2102003118

28 McGowan, S., Buckle, A. M., Mitchell, M. S., Hoopes, J. T., Gallagher, D. T., Heselpoth, R. D., et al. (2012) X-ray crystal structure of the streptococcal specific phage lysin PlyC. Proc. Natl. Acad. Sci. U.S.A. 109, 12752–12757 10.1073/pnas.1208424109

29 Xu, Q., Sudek, S., McMullan, D., Miller, M. D., Geierstanger, B., Jones, D. H., et al. (2009) Structural Basis of Murein Peptide Specificity of a γ-D-Glutamyl-L-Diamino Acid Endopeptidase. Structure 17, 303–313 10.1016/j.str.2008.12.008

30 Truong, J. Q., Panjikar, S., Shearwin-Whyatt, L., Bruning, J. B. and Shearwin, K. E. (2019) Combining random microseed matrix screening and the magic triangle for the efficient structure solution of a potential lysin from bacteriophage P68. Acta Crystallogr D Struct Biol 75, 670–681 10.1107/S2059798319009008

31 Nelson, D., Schuch, R., Chahales, P., Zhu, S. and Fischetti, V. A. (2006) PlyC: A multimeric bacteriophage lysin. Proc. Natl. Acad. Sci. U.S.A. 103, 10765–10770 10.1073/pnas.0604521103

32 Mainardi, J.-L., Legrand, R., Arthur, M., Schoot, B., Van Heijenoort, J. and Gutmann, L. (2000) Novel Mechanism of β-Lactam Resistance Due to Bypass of DD-Transpeptidation in Enterococcus faecium. Journal of Biological Chemistry 275, 16490–16496 10.1074/jbc.M909877199

33 Smith, R. E., Michno, B. J., Christena, R. L., O’Dea, F., Davis, J. L., Lidbury, I. D. E. A., et al. (2025) Enterococcal cell wall remodelling underpins pathogenesis via the release of the Enteroccocal Polysaccharide Antigen (EPA). *PLoS Pathog* (LaRock, C., ed.) 21, e1012771 10.1371/journal.ppat.1012771

34 Eckert, C., Lecerf, M., Dubost, L., Arthur, M. and Mesnage, S. (2006) Functional Analysis of AtlA, the Major *N* -Acetylglucosaminidase of *Enterococcus faecalis*. J Bacteriol 188, 8513–8519 10.1128/JB.01145-06

35 Doublié, S. (1997) [29] Preparation of selenomethionyl proteins for phase determination. In Methods in Enzymology, pp 523–530, Elsevier 10.1016/S0076-6879(97)76075-0

36 Van Den Belt, M., Gilchrist, C., Booth, T. J., Chooi, Y.-H., Medema, M. H. and Alanjary, M. (2023) CAGECAT: The CompArative GEne Cluster Analysis Toolbox for rapid search and visualisation of homologous gene clusters. BMC Bioinformatics 24, 181 10.1186/s12859-023-05311-2

37 Abramson, J., Adler, J., Dunger, J., Evans, R., Green, T., Pritzel, A., et al. (2024) Accurate structure prediction of biomolecular interactions with AlphaFold 3. Nature 630, 493–500 10.1038/s41586-024-07487-w

38 Van Kempen, M., Kim, S. S., Tumescheit, C., Mirdita, M., Lee, J., Gilchrist, C. L. M., et al. (2024) Fast and accurate protein structure search with Foldseek. Nat Biotechnol 42, 243–246 10.1038/s41587-023-01773-0

39 Meng, E. C., Goddard, T. D., Pettersen, E. F., Couch, G. S., Pearson, Z. J., Morris, J. H., et al. (2023) UCSF CHIMERAX : Tools for structure building and analysis. Protein Science 32, e4792 10.1002/pro.4792

40 McCoy, A. J., Grosse-Kunstleve, R. W., Adams, P. D., Winn, M. D., Storoni, L. C. and Read, R. J. (2007) Phaser crystallographic software. Journal of applied crystallography 40, 658–674 10.1107/S0021889807021206

41 Winn, M. D., Ballard, C. C., Cowtan, K. D., Dodson, E. J., Emsley, P., Evans, P. R., et al. (2011) Overview of the {\it CCP}4 suite and current developments. Acta Crystallographica Section D 67, 235–242 10.1107/S0907444910045749

42 Emsley, P., Lohkamp, B., Scott, W. G. and Cowtan, K. (2010) Features and development of {\it Coot}. Acta Crystallographica Section D 66, 486–501 10.1107/S0907444910007493

43 Croll, T. I. (2018) *ISOLDE* : a physically realistic environment for model building into low-resolution electron-density maps. Acta Crystallogr D Struct Biol 74, 519–530 10.1107/S2059798318002425

44 Williams, C. J., Headd, J. J., Moriarty, N. W., Prisant, M. G., Videau, L. L., Deis, L. N., et al. (2018) MolProbity: More and better reference data for improved all-atom structure validation. Protein Science 27, 293–315 10.1002/pro.3330

